# Gender differences in attitudes toward death among Chinese university students: *A survey in hunan and Heilongjiang province*

**DOI:** 10.1101/578955

**Authors:** Yuwei Wang, Siyuan Tang, Xin Hu, Chunxiang Qin, Lin Ma, Yang Li, Kaveh Khoshnood, Mei Sun

**Author notes:** **Corresponding author:** (MS).

## Abstract

**Background:** A positive attitude toward death has significant implications for college students, and can help students establish a healthy concept of life. But most colleges and universities in Mainland China have not yet carried out systematic death education courses.

**Objective:** This study aims to explore the attitudes of college age students to determine how they approach the idea of death using questions that explore five separate dimensions of attitude and belief.

**Methods:** We invited students from seven colleges in Mainland China using invitations sent to each year’s WeChat group. Students participated by scanning a QR code, and were then directed to a website that contained a self-administered questionnaire. We received 1,206 completed interviews.

**Results:** We found evidence of a substantial gender difference in attitudes toward death. These differences remain after adjustment for differences between male and female in other correlates of death attitudes, and are not a function of gender differences in the dimensionality of the five scales used to characterize attitudes.

**Conclusion:** Using previous research on gender differences as a guide, we speculate that these differences originate in culturally-defined expectations that are gender-related, as well as in substantial differences in individual family experiences of death. These speculations can take the form of testable hypotheses that should explain differences within genders as well as between genders. We believe that better education about death for college students can shape healthier attitudes among both male and female.

## Introduction

Death is the end of life. No one can avoid it. As natural as our birth, it is an inherent part of life, a natural episode of human existence [1]. Attitudes toward death vary and depend on the culture, race and gender roles that shape a person’s views of both life and death [2–4]. Death attitudes refer to people’s emotional reactions, evaluations and behavioral tendencies in response to the inevitable fact of death and ideas about death [5–8]. Initially, the measurement of attitudes toward death was mainly used as a tool to evaluate the effectiveness of death education, and focused on fear and anxiety. Subsequently, researchers have expanded this single dimension measurement to multi-dimensional measurement [9]. The current trend is to study a wide range of responses to death [10–13]. Wong, Reker and Gesser suggested that [14] research on attitudes toward death often ignores people’s natural and positive reactions. They therefore modified the concept of death attitudes to measure not only fear and avoidance, but also to include different forms of acceptance.

Several cross-cultural studies have found that men and women have different profiles of death attitudes. Chistopolskaya’s study found that [15] regardless of age, females were more likely to accept death than men. Bassett1’s study showed that [16] females scored higher than males in many fear of death items, and obtained higher scores in different dimensions of acceptance. Wong’s study reported that [14] women are less likely to avoid thinking about death and to express acceptance at higher levels than men. Power and Smith’s study showed that [17] women scored higher than men only in fearing death of people who mattered to them personally. Long pointed out that [18] cultural differences, in particular culturally-defined gender roles, may affect people’s understanding and acceptance of death.

“Death education” originated in the United States in 1928 and rose in the late 1950s. It has become very popular in developed countries [19–22]. Unfortunately, it started late in mainland China, and its popularity is low. Most colleges and universities have not yet carried out systematic death education courses [23–24]. This is also an important reason for the lack of correct attitude towards death among Chinese college students.

This study investigates the attitudes towards death and factors that influence these attitudes among students at comprehensive colleges and medical colleges in both northern and southern provinces (Hunan and Heilongjiang) in China. We document the nature of differences between men and women in our samples, and suggest a number of reasons for the differences we observed. Such differences suggest that persistent gender-defined roles in China shape comparisons between men and women. These factors affecting attitudes towards death and gender differences in attitudes towards death between male and female identify important issues that should be addressed in the death education of Chinese college students. This is a subject only recently recognized as a need in Chinese college curricula.

## Methods

### Design

A descriptive cross-sectional study was adopted for this research.

### Settings and sample

We sampled 1,254 students from seven colleges and universities - including Central South University, National University of Defense Science and Technology, Hunan Normal University, Hunan University of Traditional Chinese Medicine, Heilongjiang Qiqihar Medical College, Jiamusi University and Harbin Normal University - through a combination of cluster sampling and random sampling. After data cleaning, 1,206 questionnaires remained.

In order to identify the cultural differences between Northern and Southern China, we chose Hunan as it is located in Southern part of China, and Heilongjiang province part of Northern China. Among the seven universities and colleges, there are two medical colleges, two normal universities and three comprehensive universities. Among all participants, 536 are from Hunan province, while 670 are from Heilongjiang province. The inclusion criteria were freshman to senior students who had not yet interned or who were currently interning, and who voluntarily participated in this study.

### Ethical considerations

The research ethics committee of the Xiang Ya Nursing School of Central South University approved the study (IRB Approval Number:2018018). The participants were informed that they were taking part in the study voluntarily and anonymously. They could withdraw at any time and had the right to ignore questions they did not want to answer. Whatever they chose to do would not jeopardize their employment conditions.

### Data collection

The data were collected from April 2018 to June 2018, and 1,206 questionnaire meeting the inclusion criteria. During the investigation, the response rate to the questionnaire was 96.2%.

Data were collected with questionnaire on the network by the researcher. This software comes from Changsha Questionnaire Star Network Technology Co., Ltd. We invited students from 7 colleges in Mainland China using invitations sent to each year’s WeChat group (WeChat, like Facebook, is a platform communication tool that supports single and multi-person participation using software that sends voice, pictures, video, text, and links over the Internet). Students participated by scanning a QR code and were directed to a website that contained a self-administered questionnaire for completion.

### Data collection tools

Data were collected with a questionnaire that includes demographic information, with 21 questions related to the students’ background and their views on the issue of death, as well as the Chinese version of the Death Attitude Profile-Revised (DAP-R-C) by Zhu Hailing [25].

This revised version is adapted from the Death Attitude Profile-Revised (DAP-R) by Wong, Reker and Gesser in 1994 [11]. After translating and retranslating the original scale, according to expert consultation and pre-test feedback results, on the basis of ensuring the equivalence of the scale, ambiguous items 4, 18 and 22, and repetitive item 20 were deleted, and items 8, 16, 25, and 31, which had similar meanings, were merged to determine the content of the DAP-R-C scale. The original 32 items were reduced to 25 items. These adapted questions were fewer in number and easier to complete.

The Cronbach’s α of each DAP-R-C subscale was between 0.585~0.853, and the correlation coefficients between the subscales are smaller than the Cronbach’s α of each subscale. The Cronbach’s α of the total scale was 0.840. The split-half reliability of each DAP-R-C scale was between 0.598~0.809. The total scale was divided into two equal parts by odd and even half method, and the split-half reliability of the total scale was 0.843. In this research, 20 college students participated in the pilot study and found the questions were understandable and appropriate according to the research purpose.

The scale measures five dimensions of attitudes toward death: Approach acceptance, Escape acceptance, Fear of death, Death Avoidance, and Neutral acceptance.

1. Approach acceptance: Individuals believe there will be a better after-life after death. They regard death as a channel to happiness. They easily accept the concept of death, and even hope that death will come sooner rather than later.
2. Escape acceptance: Individuals fear life more than death. They regard death as a way to relieve the pain of life. It is a kind of death acceptance forced by suffering.
3. Fear of death: Refers to the individual’s fear and negative emotions when facing death.
4. Death Avoidance: Individuals try their best to avoid thinking about death or discussing things related to death, and consider talking about death to be taboo.
5. Neutral acceptance: Individuals believe that death is part of the process of life. It is natural and unavoidable. Those who hold such attitudes do not fear or welcome death. They regard death only as a natural stage of life.

Each subscale has five items, using a Likert 5-point scale scoring method, that is, “strongly disagree” scoring 1 point; “disagree” scoring 2 points; “neither agree nor disagree” scoring 3 points; “agree” scoring 4 points; “strongly agree” scoring 5 points.

## Results and findings

Among a total of 1,206 college students, including 388 males and 818 females, the highest score was found in the dimension of neutral acceptance and the lowest in the dimensions of approaching acceptance and escape acceptance. Gender is the only characteristic associated with all five dimensions of death attitude measures in our study. Table 1 shows the average scores for the full sample and for men and women separately. College students had the highest total scores on the neutral acceptance dimension. Female students were significantly higher than male students on all of the neutral acceptance items and with respect to total natural acceptance scores. In comparison, men were significantly higher than women in the dimensions of fear of death, death avoidance, approach acceptance and escape acceptance.

**Table 1.** Average Scores on Five Death Attitude Scales^#^ of Chinese College Students

| Death attitude | Totality<br>( n=1206 ) | Female students<br>( n=818 ) | Male students<br>( n=388 ) | t | p |
| --- | --- | --- | --- | --- | --- |
| Neutral Acceptance | 19.02±3.97 | 19.39±3.80 | 18.25±4.21 | 4.52 | 0.001** |
| Approach Acceptance | 11.33±4.50 | 11.04±4.47 | 11.93±4.51 | -3.33 | 0.001** |
| Escape Acceptance | 12.64±3.85 | 12.36±3.81 | 13.24±3.87 | -3.74 | 0.001** |
| Fear of Death | 13.43±4.27 | 13.18±4.31 | 13.96±4.15 | -2.98 | 0.003 |
| Death Avoidance | 12.57±4.61 | 12.09±4.59 | 13.57±4.48 | -5.3 | 0.001** |

\*\*p<0.001 <sup>#</sup> DAP-R-C=Chinese version of the Death Attitude Profile-Revised.

Table 2 shows the correlations between the five subscales. In general, these correlations are larger than the ones observed in the study of Hong Kong students [26], but are nearly always of the same sign (direction). In our study, the four subscales that show men’s averages higher than women’s averages have fairly large and significant positive correlations. On the other hand, Neutral Acceptance, the only subscale where women’s average scores exceed those of men, shows negative correlations with all of the other subscales.

**Table 2.**
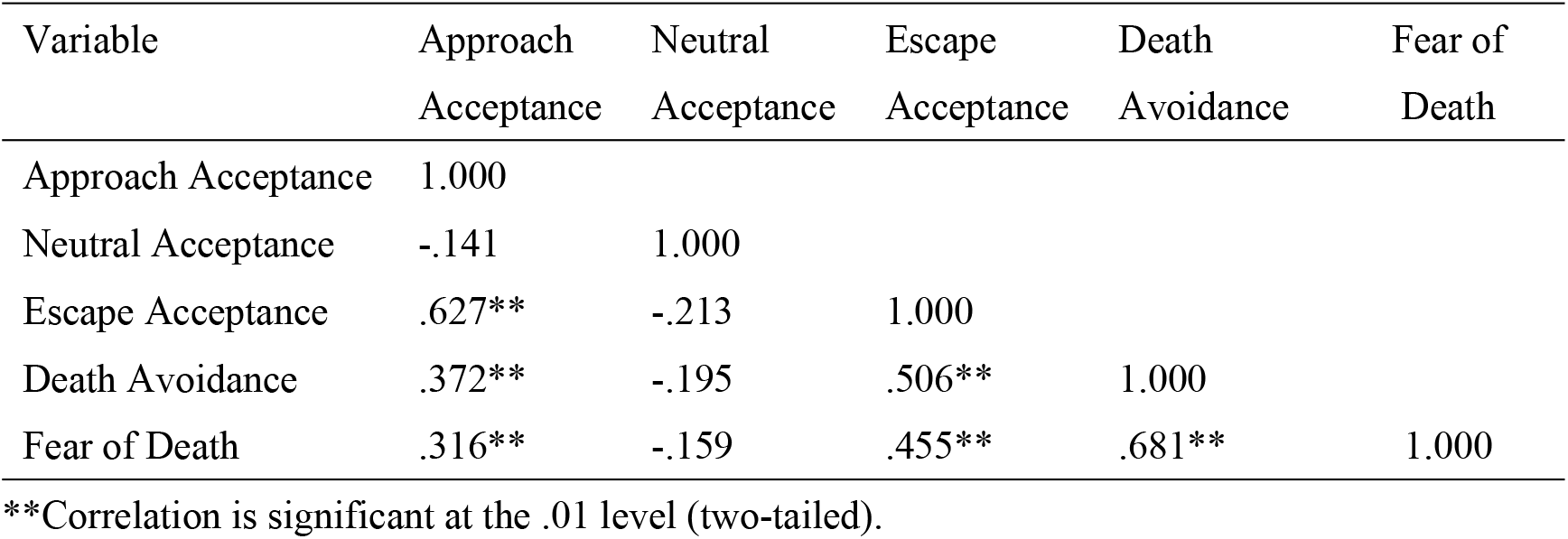
Correlations Between Students’ Responses to the Five Dimensions of DAP-R-C

In order to determine whether the dimensions characterizing males and females are the same or different, we examine the correlations separately for each gender. These are shown in Table 3 below. The magnitudes and the signs of the correlations are very similar in males and females. This suggests that the dimensions underlying the subscales are actually the same. The major difference is where males and females are located on the scales. So the average differences tell an even stronger story about gender differences, because males and females can be placed on the same subscales rather than in gender-specific dimensions.

**Table 3.**
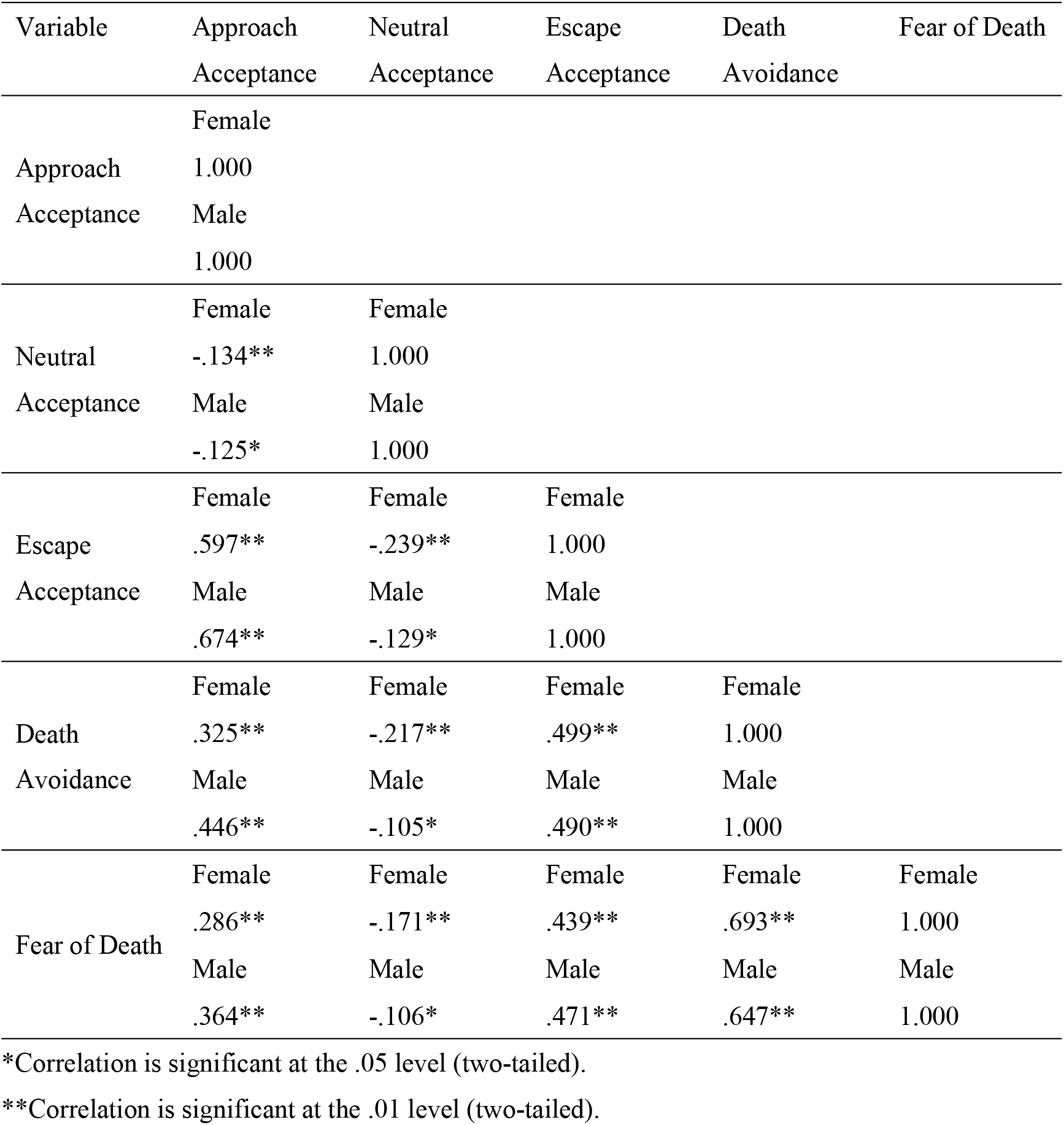
Correlations Between Each Gender’s Responses in Five Dimensions

To investigate why men and women have such a different profile of attitudes about death, we examined other factors in the data that distinguish men from women in this sample. Those that show significant differences between men and women are presented in Table 4 below.

**Table 4.** Factors Distinguishing Men and Women in Our Sample of College Students

| Factor | P-value for male<br>versus female | Direction of<br>difference |
| --- | --- | --- |
| Single child | P<.000 | Female > Male |
| Academic performance | P<.008 | Female > Male |
| Past or current serious illness | P<.003 | Male > Female |
| Past or current mental illness | P<.006 | Male > Female |
| Family discusses death openly | P<.011 | Female > Male |
| Medical versus non-medical discipline | P<.000 | Female > Male |
| College Year | P<.000 | Female > Male |

\*P<0.05

Table 5 shows that the factors distinguishing men and women in our sample do not explain the persistent gender differences in attitudes toward death. Gender is a highly significant predictor of all death attitudes controlling for all differences between genders listed in Table 5. Year in school, a surrogate for age differences in this relatively age-homogeneous sample, mental illness and discussions about death in the home are factors that have significant associations with most of the five scales. However, gender differences persist when adjusting for these gender-differentiated variables. None of the differences we observe in Table 4, even those that appear as significant predictors of death attitudes in Table 5, can explain the gender difference.

**Table 5:** Regression of Death Attitude Factors on Gender and Factors Related to Gender in Our Sample

| Predictive Factors | Approach<br>Acceptance | Neutral<br>Acceptance | Escape<br>Acceptance | Death<br>Avoidance | Fear of<br>Death |
| --- | --- | --- | --- | --- | --- |
| Gender F=1, M=2 | 0.868*** | -0.981**** | 0.961**** | 1.357**** | 0.836*** |
| Single child =1,<br>Not Only=2 | -0.497* | 0.099 | -0.359 | -0.176 | -0.161 |
| Grade = 1,2, or 3 | -0.331** | 0.296** | -0.097 | -0.361** | -.334** |
| Had or has serious<br>illness=1; No=2 | 0.309 | 0.339 | 0.005 | -0.115 | -0.293 |
| Had or has mental<br>illness=1; No=2 | -0.962* | 0.912** | -1.988**** | 0.405 | 0.055 |
| Talk about death in<br>family: 1=yes, | 0.134 | -0.363* | 0.337* | 1.095**** | 0.787**** |
| 2=when necessary,<br>3=never |  |  |  |  |  |
| Major: Medical=1,<br>Non-Medical=2 | -0.175 | -.006 | 0.235 | 0.211 | -0.143 |
| Grade performance:<br>1=excellent,2=good<br>3=fair,4=poor | 0.065 | 0.312** | -0.250 | -0.285* | 0.221 |
| Multiple Correlation<br>Coefficient | .141 | .184 | .191 | .223 | .161 |

\*\*\*\*p<.001, \*\*\*p<.01 \*\*p<.05, \*p<.10

Table 6 shows that the largest gender differences appear on several items that provoke substantial disagreement from both genders. Over these items target avoidance of thinking about death, the concept that death ends everything, and the idea that death is an escape from suffering. Although both men and women frequently reject these ideas, women show a stronger rejection. The average gender differences in subscales reflect these items more than others. While many respondents of both genders do not shy away from thoughts of death, women appear more likely to allow these thoughts into their consciousness. Furthermore, most women do not their see their own death as the end of everything, perhaps because their societal role emphasizes the preparation of a generation that will survive them. Nearly two-thirds of women rejected the idea that death is an escape from suffering, compared to only half of the men. These results suggest that the gender differences we observe are not of a piece. They appear for some kinds of attitudes and disappear for others. We see that most women in our sample do not push death out of their thoughts; they “do” believe in something beyond death, and are not inclined to see death as a way to avoid suffering. Many men also feel this way, but they don’t show the higher level of rejection that we see in women.

**Table 6:**
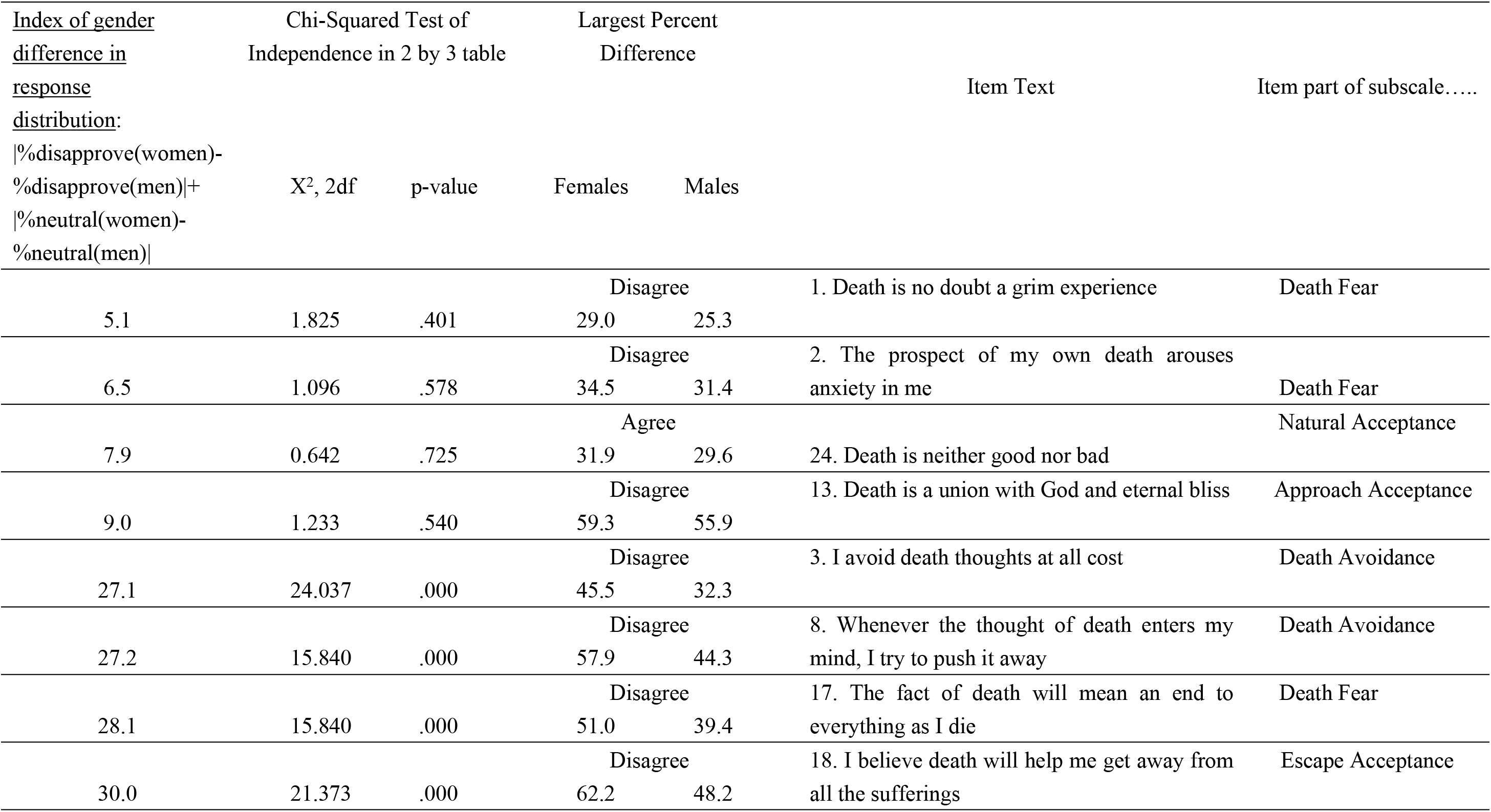
Items with the largest and smallest gender differences in response distributions

## Discussion

Many aspects of traditional China that distinguish men and women continue to powerfully influence gender roles, even though China has entered an unprecedented period of cultural and social modernization. The rapid improvement in women’s social status is important for family behavior in China. Women no longer live under traditional Chinese patriarchal rules, that is, obeying their father at home and obeying their husband after marriage [27]. Since the founding of the People’s Republic of China, women’s social and economic status has improved significantly, but women’s economic status is still lower than men’s. Today, women have increasingly free speech, in particular since 1950 [28], when China promulgated the Marriage Law, which formally legalized women’s free speech and equalized the rights and interests of wives and husbands [29]. Although women’s social and family status has to some extent improved, men still take on greater social and family responsibilities, due to the influence of traditional Chinese ideology and culture [30]. In China, male university students focus on going out into society after graduation, with the need to achieve in their career and to support a family [31]. Soaring property prices in China will undoubtedly add considerable burden to their future lives. Below, we consider some of the reasons we observe gender differences in attitudes toward death, keeping in mind the overall context of culturally-driven gender roles that persist, even in a society that is modernizing rapidly.

### Possible explanations for our findings on gender differences

#### a. Women’s Greater Role in Preparing for the Next Generation and Greater Experience in Caring for the Elderly

Women will have many roles in the family after marriage [32]: often taking care of two sets of aging parents, managing relations between relatives and raising children. Chinese female college students have typically not yet experienced these things, but when they occur, these changes can exert a profound effect on their lives. In the process of living their lives and taking on new roles, women experience birth as they become a mother for the first time, and the continuity of the life process. At the same time, since women are on the verge of life and death during childbirth, they also experience the approach of death.

A mother’s care is an indispensable growth experience for each person from childhood onward. The renowned education expert Froebel once said: “The fate of the nation is not so much in the hands of the authorities as in the hands of the mother.” This point of view profoundly illustrates the important role of women in fostering the next generation [33]. From the perspective of the origin of roles, women naturally acquire the identity of mother from the moment a child is born. This identity is based on blood relationship and is accompanied by a child’s birth. Therefore, the influence of the mother on the child is the child’s earliest influence [34]. Chinese women regard their children as a continuation of their own lives, which means they have a huge investment in the future beyond their own lives, so they are not afraid that death will end everything.

Due to their traditional role taking care of elderly parents [35], women already have the experience of preparing for death; they also have psychological endurance. The coexistence of evidence and reality shows that women tolerate pain more easily and have a greater acceptance of death than men. Female students in China, especially adults, look forward to experiencing the glory of motherhood. Mothers love their children and are devoted to their families. Females, therefore, tend to have more communication with their mothers than with their fathers, and they have the freedom to speak with their mothers about death. This is influenced by many factors, such as the mother’s management of the family and her caring for the elderly, to them, death is more acceptable. Females must face the reality that a couple will support four old people and raise one or two children in the future. This is the result of China’s special national conditions. In the past, in order to control population growth, China adopted an “only child” policy, that is, couples were permitted to have only one child. In recent years, however, China liberalized that policy, replacing it with a “two-child policy”, that is, after graduation from college couples may now have one child or more. Today, married couples now need to support two sets of parents and raise at least one child of their own.

#### b. Men have more stress in their role as family providers and often don’t talk about their anxieties and fears

As Chinese boys grow up, their masculine identity continues to develop. Influenced by their father as a male role model, their focus has always been on how men should shoulder the burdens of life and bring happiness and security to their family [36]. Men talk about their dreams, their future and their responsibilities, but seldom discuss life and death with their children. In addition, they believe that it is too heavy, even unlucky, to talk to a child about death. Even if the child talks about death, fathers tend to avoid the topic. In a culture where marriage is an expected part of adult life and with a deepening of economic pressures, it becomes more difficult for men to start and maintain a marriage. In traditional Chinese culture, men have a sense of superiority [37]. The traditional Chinese concept of maleness emphasizes that men’s work centers around the outside world and that men must shoulder the responsibility for all outside difficulties [27].

## Conclusion and implications

### Implications for death education

Professor Hu of Guangzhou University once said that “Death education is not to beautify death, but to remove the mystery of death and to illuminate the sacredness of death, so that students cherish life more, and enhance their psychological health [38].” At the same time, we must formulate China’s death education policy in light of China’s special national conditions. Death education should be compulsory in all university classes in China (including medical schools), to avoid the fear of death and an escapist attitude among young people. Increasing the availability of psychological counseling for boys, in particular, can help them avoid the stress of excessive pressures in life and prevent them from developing an inaccurate view of death. Students should be encouraged to talk more about life and death, discuss this topic with their parents when appropriate, and cooperate with and guide their own children to establish a correct concept of death. They should be encouraged to avoid feeling embarrassed about being an only child unwilling to seek psychological assistance. As only children, many Chinese had no siblings to share their thoughts and feelings with, and may be afraid of their parents knowing their secrets. When students are educated about death, as they grow older, they can understand the meaning of life and death more profoundly, thus allowing them to cherish their learning environment and opportunities, correct their learning attitudes, lay a foundation to improve their academic performance, and play a positive role in preventing the occurrence of suicide and suicide attempts – whether for themselves or for others - in the future.

### Topics for further research

Our interpretations lead to research questions that can be investigated in future research focusing on college students and older adults alike. Here are three, all of which involve direct measurement of the factors mentioned above for individual men and women.

1. Are women who are involved in childrearing less likely to see death as an end to everything than women with no children?
2. Are women who care for elderly parents more likely to think about death and talk openly about death than women who don’t act as caretakers?
3. Are men who show higher levels of stress about their job and finances more likely to see death as an escape from life than men who are less stressed?

## Acknowledgments

We thank Dr James A. Wiley for data validation and all the study participants.

## Author Contributions

Conceptualization: Mei Sun

Investigation: Yuwei Wang, Xin Hu

Methodology: Chunxiang Qin, Kaveh Khoshnood

Project administration: Siyuan Tang

Resources: Lin Ma, Yang Li

Visualization: Siyuan Tang

Writing – original draft: Yuwei Wang

Writing – review & editing: Kaveh Khoshnood, Mei Sun

